# Biofluid modeling of the coupled eye-brain system and insights into simulated microgravity conditions

**DOI:** 10.1101/609958

**Authors:** Fabrizia Salerni, Rodolfo Repetto, Alon Harris, Peter Pinsky, Christophe Prud’homme, Marcela Szopos, Giovanna Guidoboni

**Affiliations:** Department of Mathematical, Physical and Computer Sciences, University of Parma, Parma, Italy; Department of Civil, Chemical and Environmental Engineering, University of Genoa, Genoa, Italy; Eugene and Marilyn Glick Eye Institute and Department of Ophthalmology, Indiana University School of Medicine, Indianapolis, IN, USA; Department of Mechanical Engineering, Stanford University, Stanford, CA, USA; Institute of Advanced Mathematical Research UMR 7501, University of Strasbourg CNRS, Strasbourg, France; Laboratoire MAP5 (UMR CNRS 8145), Université Paris Descartes, Sorbonne Paris Cité, France; Department of Electrical Engineering and Computer Science, Department of Mathematics, University of Missouri, Columbia, MO, USA

## Abstract

This work aims at investigating the interactions between the flow of fluids in the eyes and the brain and their potential implications in the development of visual impairment in astronauts, a condition also known as spaceflight associated neuro-ocular syndrome (SANS). To this end, we propose a reduced (0-dimensional) mathematical model of fluid flow in the eyes and brain, which is embedded into a simplified whole-body circulation model. In particular, the model accounts for: *(i)* the flows of blood and aqueous humor in the eyes; *(ii)* the flows of blood, cerebrospinal fluid and interstitial fluid in the brain; and *(iii)* their interactions. The model is used to simulate variations in intraocular pressure, intracranial pressure and blood flow due to microgravity conditions, which are thought to be critical factors in SANS. Specifically, the model predicts that both intracranial and intraocular pressures increase in microgravity, even though their respective trends may be different. In such conditions, ocular blood flow is predicted to decrease in the choroid and ciliary body circulations, whereas retinal circulation is found to be less susceptible to microgravity-induced alterations, owing to a purely mechanical component in perfusion control associated with the venous segments. These findings indicate that the particular anatomical architecture of venous drainage in the retina may be one of the reasons why most of the SANS alterations are not observed in the retina but, rather, in other vascular beds, particularly the choroid. Thus, clinical assessment of ocular venous function may be considered as a determinant SANS factor, for which astronauts could be screened on earth and in-flight.

## Introduction

Microgravity conditions have been observed to induce visual function alterations in many astronauts that pose serious challenges for both astronauts and their missions in space [1, 2]. This syndrome, also known as spaceflight associated neuro-ocular syndrome (SANS), is characterized by a large number of apparently unrelated and often not concurrent symptoms. These include choroidal folds, cotton wool spots, optic nerve distension and/or kinking, optic disc protrusion, posterior globe flattening, refractive deficits and elevated intracranial pressure [3]. Added to the complexity of the range of symptoms are the problems of susceptibility and genetic predisposition to develop visual problems.

The current understanding of how weightlessness environment affects the human body and may lead to SANS development is still quite rudimentary. Various studies of the symptoms experienced by astronauts during long-term missions (four to six months) have been performed [1], but their validity is hampered by the small size of the subjects cohort. To overcome this difficulty, ground-based microgravity laboratory models have been proposed, the most significant of which is the long head down tilt (LHDT) experimental procedure that is used to simulate the effects of microgravity on the cardiovascular system. LHDT experiments have shown that the fluid shift caused by the tilt produces a transient increase in central venous pressure, later followed by an increase in left ventricular size without changes in cardiac output, arterial pressure, or contractile state [4]. Moreover, experiments performed by [5] confirmed that microvascular pressures measured in the lower lip increase, as well as subcutaneous and intramuscular interstitial fluid pressures in the neck. Interestingly, interstitial fluid colloid osmotic pressures remain unchanged, whereas plasma colloid osmotic pressures drop significantly after 4 hours of LHDT, thereby suggesting a transition from fluid filtration to absorption in capillary beds between the heart and feet during LHDT. The above-mentioned studies suggest that two main mechanisms may be involved in SANS pathophysiology, namely: *(i)* changes in the vascular system and fluid distribution (fluid shifts); and *(ii)* changes in the brain/central nervous system and intracranial pressure (ICP). However, many factors may influence these changes, including blood pressure, intraocular pressure (IOP) and cerebrospinal fluid pressure (CSFp), which are difficult to isolate in an experimental setting.

A complementary approach to experimental studies is the use of mathematical modeling as a tool to investigate theoretically the role of various factors potentially contributing to SANS and help elucidate the mechanisms of their interactions and their implications in the loss of visual function. To the best of our knowledge, the mathematical modeling of hemo-fluid dynamic contributions to SANS is still at its early stages. A lumped-parameter model to simulate volume/pressure alterations in the eye in zero gravity conditions was presented in [6]. The model includes the effects of blood and aqueous humor dynamics, ICP, and IOP-dependent ocular compliance. A lumped-parameter model of fluid transport in the central nervous system aimed at simulating the influence of microgravity on ICP was presented in [7]. The model could be coupled, in the authors’ intention, to lumped parameter and finite element models of the eye. The present work advances the modeling methods in the context of SANS by providing a more detailed description of fluid circulations in the eyes and their connections to the brain, as sketched in [8–10]. In particular, the model accounts for the blood flow within three major ocular vascular beds, namely retina, choroid and ciliary body. The model describes the intimate connection between ocular and cerebral hemodynamics, as brain and eyes share vessels for blood supply and drainage. In addition, the cerebrospinal fluid (CSF) flowing within the subarachnoid space influences the tissue pressure in the optic nerve head and, as a consequence, the biomechanics and hemodynamics of the lamina cribrosa and the vessels running through it. The model is used to investigate the complex relationship between blood pressure, IOP, ICP, CSFp and the flow of fluids within eyes and brain, which are thought to be important factors contributing to SANS.

Interestingly, the modeling ingredients embedded in the proposed model make it a very flexible tool to study interactions between the eyes and the rest of the body, particularly the brain, in various conditions beyond the microgravity environment. Thus, besides the application to SANS discussed in the present work, we believe that the proposed model could serve as a basic framework to study pathological states, most importantly glaucoma [11–13], where the interaction between ocular and cerebral hemo-fluid-dynamics is thought to play an important role. Specifically, it is well-documented that the levels of blood pressure, IOP and CSFp, along with other vascular and biomechanical risk factors, strongly affect etiology, progression and incidence of glaucoma [14–18]. As medicine is clearly heading in the direction of tailored diagnosis and treatment, the clinical ability to ascertain the relative weight of each risk factor in any given patient is of paramount significance. Although the use of imaging technology in medicine has advanced in giant steps in recent years, there is no clinical standard for quantifying all relevant clinical outcomes in any given subject. In this perspective, mathematical modeling can provide a virtual lab where the clinical significance of each risk factor can be quantified and novel individualized therapeutic approaches can be designed [19].

The paper is organized as follows. We begin by providing a description of the mathematical model, which is articulated in four main parts, first outlining its general characteristics, then describing in more detail the model of the brain, of the eyes and of their coupling. We then discuss how microgravity conditions are modeled and how the overall system is solved numerically. A description of the rationale adopted for model calibration is followed by an outline of the model validation against experimental observations. The paper ends with the results of model simulations and their critical discussion.

## Materials and methods

### Lumped parameter mathematical model

#### General description of the model

We propose a lumped parameter model of the brain and eyes, connected with a highly simplified model of the body, the electrical analogue of which is shown in Fig 1. The aim of the model, which describes pressure and flow distributions in the brain and eyes, is to predict fluid redistribution in the upper body vasculature and variations of the IOP and ICP following exposure to a microgravity environment.

**Fig 1.**
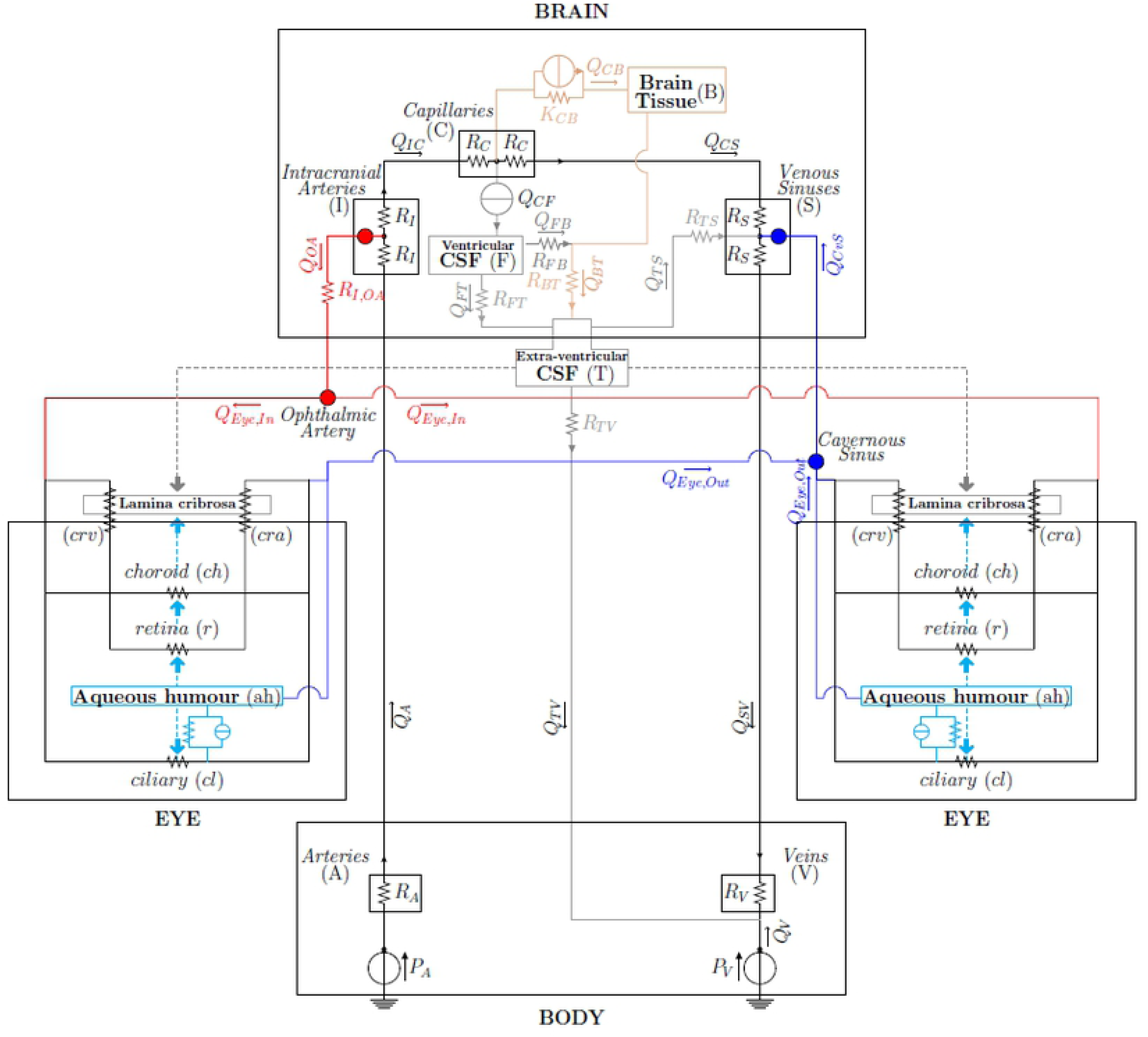
Network model of fluid flows in the brain and eyes. The nodes correspond to the connection between the brain and eye models. The connection *Intracranial Arteries-Ophthalmic Artery* represents arterial supply; the connection *Venous Sinuses-Cavernous Sinus* represents the venous drainage; the grey and cyan arrows represent the pressures acting on both sides of the lamina cribrosa.

The physiological system is subdivided into a number of linked, interacting compartments, each of which contains a single physical constituent, such as blood, cerebrospinal fluid, interstitial fluid and aqueous humor. In particular, we consider the interactions among the following components (see Fig 1):

##### in the brain

- blood (3 compartments: intracranial arteries (I), capillaries (C), intracranial venous sinuses (S));
- cerebrospinal fluid (1 compartment: ventricular CSF (F));
- cerebral tissue and interstitial fluid (1 compartment: brain (B));

##### in each of the two eyes

- blood (5 compartments: retina (*r*), choroid (*ch*), central retinal artery (*cra*), central retinal vein (*crv*), ciliary body circulation (*cl*));
- aqueous humor (*ah*) (1 compartment: anterior and posterior chamber);

##### and in the eye-brain coupling

- cerebrospinal fluid (1 compartment: extra-ventricular (T) bridging intracranial and extracranial regions, also including the subarachnoid space in the optic nerve posterior to the lamina cribrosa).

The lumped parameter circuit for the brain adopted in the present work was described and validated by Lakin and Stevens in [20] for microgravity simulations. The eye block combines a model for the retinal circulation and a model for the dynamics of aqueous humor that have been proposed by some of the authors of this paper in [21] and [22], respectively, and extends them to account also for the choroidal and ciliary body vascular beds [23]. In this work, we focus on the mean behavior of the system and we neglect time variations on both the short time scale of the heart beat and on the long time scale of remodelling processes. We also neglect autoregulation mechanisms of small vessels, since we wish to keep the model relatively simple in order to understand its basic behaviour. Moreover, little information exists about autoregulation mechanisms in orbit, except that they might be altered due to high *CO*_2_ concentrations [1].

The brain and eye models are coupled to each other and are linked to the rest of the body via a highly simplified model consisting of two compartments: central arteries (A) and central veins (V). This description corresponds to a model for the human physiology of the upper part of the body. Throughout the paper, lower case letters will denote compartments in the eyes and upper case letters will denote compartments in the brain and body.

Two main types of flows are included in the model, namely *filtration* and *pressure-driven flows*. Fluid filtration is accounted for in two instances: *(i)* filtration of interstitial fluid from the capillaries to the interstitial space, with the associated flow denoted by *Q*_*CB*_, and *(ii)* filtration of aqueous humor from the ciliary body capillaries into the posterior/anterior chambers, with the associated flow denoted by *J*_*uf*_. In general terms, the flux *Q*_*ij*_ due to filtration from the compartment *i* to the compartment *j* is modeled by the Starling-Landis equation [24]:

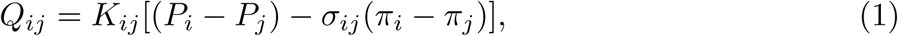

where *P*_*i*_ and *π*_*i*_ are the hydraulic and osmotic pressures in the compartment *i*, whereas *K*_*ij*_ and *σ*_*ij*_ are the filtration and reflection coefficients from the compartment *i* to the compartment *j*. The electric analogue of the Starling-Landis Eq (1) is an element with a resistor and a current generator arranged in parallel, as depicted in Fig 1. All other flows in the model are simply driven by the hydraulic pressure difference between compartments. In this case, the pressure-driven flux *Q*_*ij*_ between the generic compartments *i* and *j* is governed by the hydraulic analogue of Ohm’s law

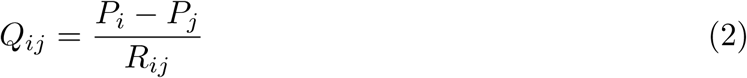

where *R*_*ij*_ denotes the hydraulic resistance. Each compartment *i* in the model is assumed to have a total resistance to flow of 2*R*_*i*_; then the resistance *R*_*ij*_ in Eq (2) is taken to be *R*_*ij*_ = *R*_*i*_ + *R*_*j*_, i.e. the sum of half of the resistance of compartments *i* and *j.* In the following, owing to the complexity of the eye model, we will use different notations to identify the various components of the eye circuits.

By formulating the above equations for all compartments and by writing the Kirchhoff law of currents at all circuit nodes, we obtain a set of nonlinear algebraic equations in the unknowns *P*_*i*_, as reported in Tables 1 and 2. The nonlinearity is a consequence of the fact that, in some compartments, resistances are assumed to depend on pressures. This is discussed in the following sections, where the brain and eye models are described in more detail.

**Table 1.**
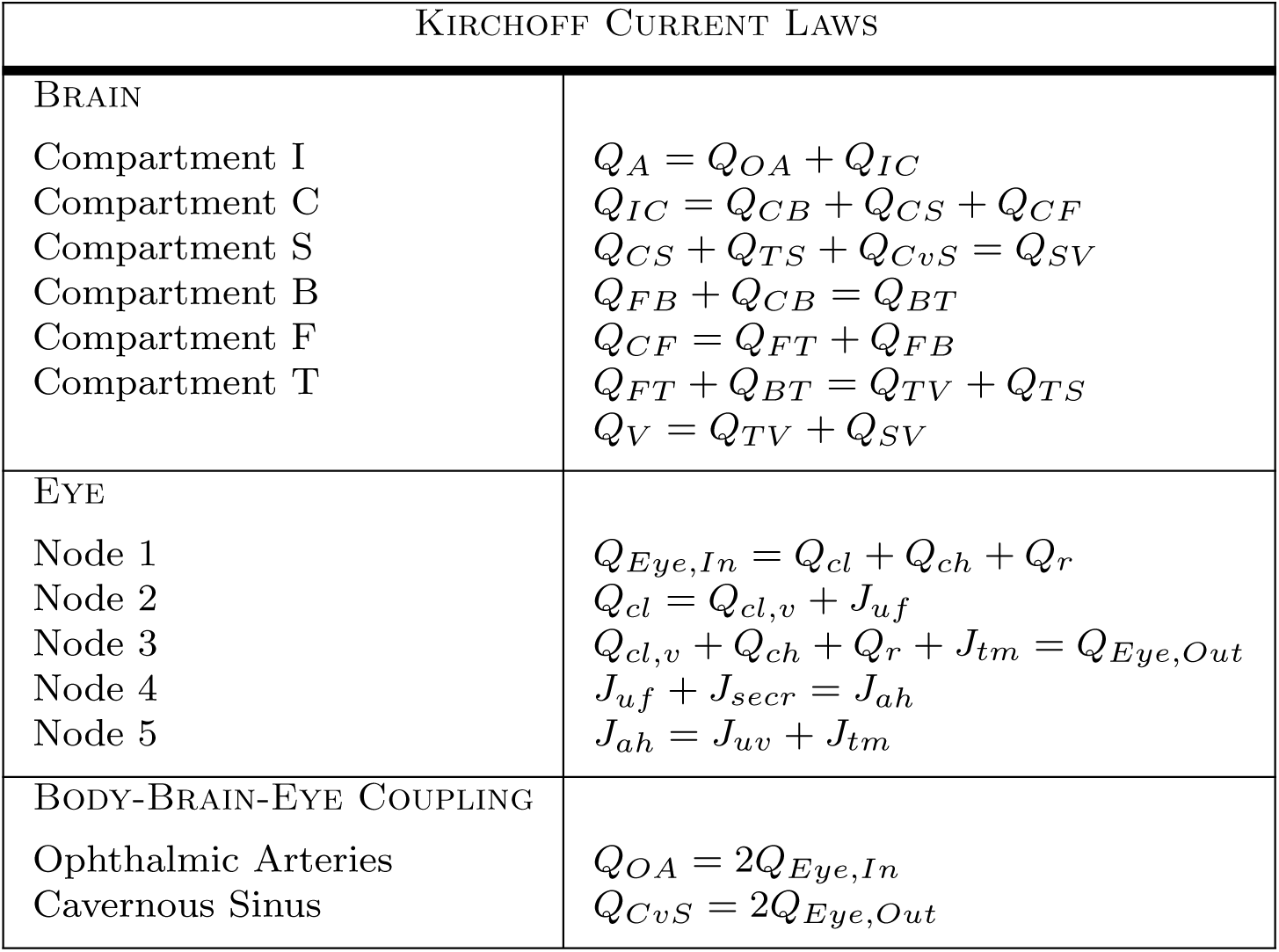
Summary of the model equations obtained by writing the Kirchhoff current law at the circuit nodes.

**Table 2.**
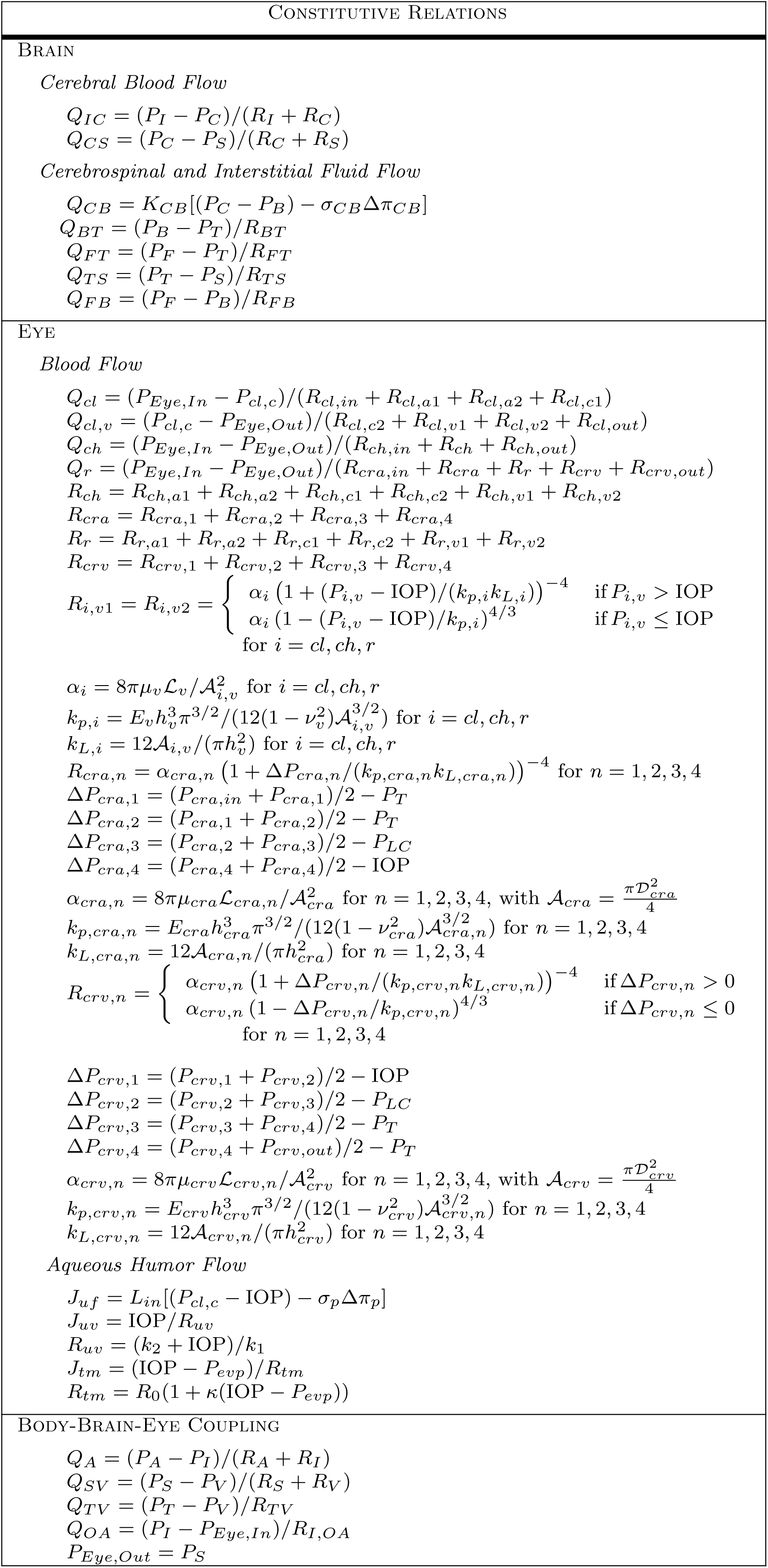
Summary of the model balance and constitutive relations.

### The brain model

The human brain is a fully enclosed organ and its tissue is a medium saturated by three different fluids interacting with each other: *(i)* blood, *(ii)* cerebrospinal fluid and *(iii)* interstitial fluid. The cerebrospinal fluid is produced mainly by the choroid plexus. It flows through the ventricular system to the subarachnoid space, where it is absorbed into the blood stream via the sagittal sinus. The interstitial or extracellular fluid fills the interstices of the brain tissue, bathing the neurons. In healthy subjects, only a relatively small amount of fluid is able to leak from blood into the interstitial fluid, owing to the low permeability of the blood-brain barrier [25].

In this work, we describe the brain via an electrical analogue representation of the model proposed by Lakin and Stevens in [20], as schematically shown in the “BRAIN” box of Fig 1. The model consists of three fluid networks: *(i)* the blood vasculature (Fig 1, black portion), *(ii)* the cerebrospinal fluid network (Fig 1, grey portion) and *(iii)* the interstitial fluid network (Fig 1, tan portion). The brain model is connected to the upper part of the body through the systemic central arteries (A) and veins (V) compartments characterized by the resistances *R*_*A*_ and *R*_*V*_, respectively. CSF production is externally imposed and kept constant at the production rate *Q*_*CF*_. Unlike the choice adopted in [20] to impose a constant flux *Q*_*IC*_ between the intracranial arteries and the capillaries accounting for the autoregulation of cerebral blood flow, here we allow cerebral blood flow to change by inserting the hydraulic resistances *R*_*I*_ and *R*_*C*_ between the compartments I and C. An extra-ventricular compartment (T) filled with cerebrospinal fluid bridges the intracranial and extracranial regions and includes the subarachnoid space in the optic nerve, posterior to the lamina cribrosa.

### The eye model

The full model of the eye is schematically shown in the two “EYE” boxes of Fig 1 and is depicted in detail in Fig 2. It comprises two interconnected circuits hosting the flow of blood and aqueous humor, respectively. The vascular circuit (Fig 2, black portion) consists of three vascular beds, corresponding to the retinal, choroid and ciliary body circulations, arranged in parallel to each other. In the aqueous humor circuit (Fig 2, cyan portion) fluid is produced in the ciliary body via ultrafiltration and active secretion, and then drained via the trabecular and uveoscleral pathways. Inclusion of this circuit is important since the balance between aqueous production and drainage governs the levels of IOP. The light blue-shaded area in the figure represents the interior of the eye globe, where the pressure equals IOP. This means that all vessels in the circuits exposed to such an area are subjected to an external pressure equal to IOP. Finally, the grey-shaded area represents the lamina cribrosa, which is loaded by IOP on the one side and by the pressure within the optic nerve tissue on the other side, which is mainly due to the CSFp in the subarachnoid space [26]. The central retinal artery and vein running through the lamina are subjected to external mechanical actions that are produced by the deformation of the lamina [27, 28].

**Fig 2.**
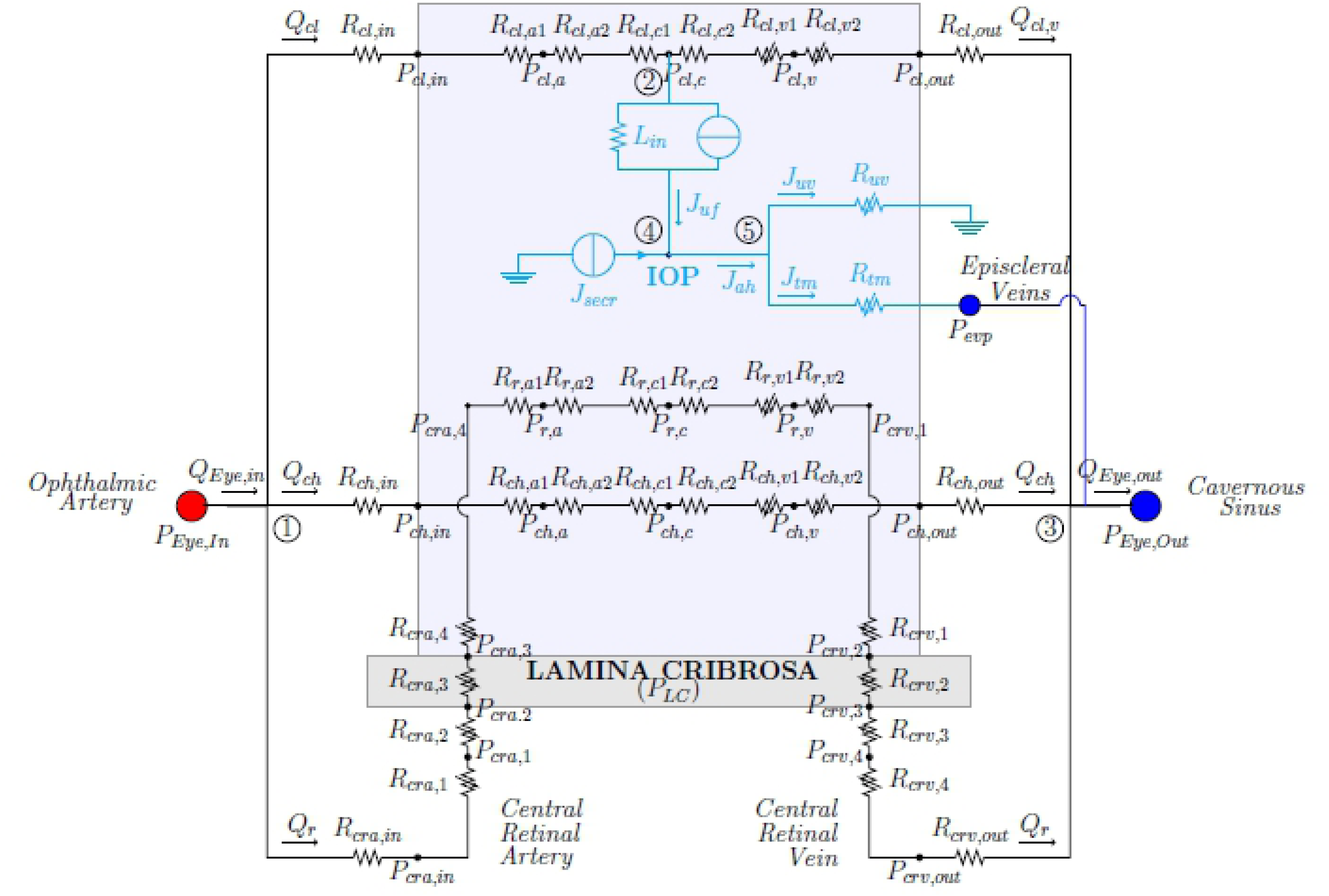
Network model for the eye vasculature (black portion) and aqueous humor production and drainage (cyan portion). The vasculature comprises circulation of blood in retinal, ciliary and choroidal vascular bed. Variable resistances are marked with arrows.

Further details of all eye circuits and the eye-brain coupling are provided in the following subsection.

#### The ocular vascular circuit

The main arterial supply to the eye is the ophthalmic artery, which branches from the carotid artery. The major branches of the ophthalmic artery are the long ciliary arteries, that nourish the ciliary body, the central retinal artery (*cra*) that nourishes the inner part of retina and the posterior ciliary arteries that nourish the choroid. Ciliary body and choroidal circulations drain into the superior ophthalmic vein, which exits the orbit through the superior orbital fissure and drains into the cavernous sinus. Retinal blood drainage occurs via the central retinal vein (*crv*) and eventually converges into the cavernous sinus. The *cra* and *crv* run through the optic nerve canal, piercing the lamina cribrosa approximately in its center.

Each vascular bed of the circuit is composed of three lumped compartments connected in series and representing arterioles (*a*), capillaries (*c*) and venules (*v*), as detailed in Fig 2. In the figure and the following text, the pressure in each node of the vascular circuit is denoted *P*_*i,n*_, where *i* = *r, cl, ch, cra, crv* indicates the different vascular segments (*r* for retina, *cl* for ciliary body, *ch* for choroid and *cra* and *crv* for the central retinal artery and vein, respectively) and *n* allows us to distinguish between different nodes within the same segment (*n* = 1, 2, 3 in the *cra* and *crv, n* = *a, c, v* in the retinal, ciliary body and choroidal circulations, and *n* = *in, out* in the vascular structures located before the arterioles and after the venules of the vascular segment *i*). A similar notation holds for all resistances; for instance, *R*_*r,v*2_ denotes the resistance in the retinal circuit downstream of the node representative of retinal venule (with pressure *P*_*r,v*_).

The vascular circuits include both constant and variable resistances. Constant resistances are calibrated at the reference state and their value is kept constant in all numerical simulations. In particular, constant resistances are adopted for arterioles and capillaries, since active changes in vascular diameters due to blood flow regulation are not accounted for in the present model. To characterize constant resistances, we used Poiseuille law to relate flux and pressure drop along each single vessel. This leads to the following expression for a generic resistance of each component of the circuit

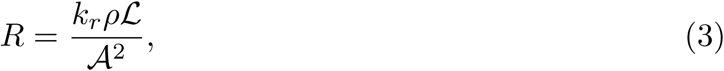

where *𝒜* is the representative cross-sectional area of the compartment, *ℒ* the representative compartment length, *ρ* is blood density, *µ* is blood dynamic viscosity and *k*_*r*_ = 8*πµ/ρ*. In the case of a single vessel, *𝒜* and *ℒ* correspond to the vessel cross-sectional area and length respectively. Also note that the circuit has been constructed in such a way that the resistance of each compartment within the eye (arterioles, capillaries and venules) is split into two resistances with the same characteristics (see Fig 2), resulting in a node representing the mean pressure of the compartment.

Variable resistances are indicated by an arrow in Fig 2. They account for passive changes in vascular diameter and, possibly, shape of the cross-sectional area, due to the action of the transmural pressure difference Δ*p*_*t*_ (i.e. the pressure difference across the vessel wall) and the deformability of the vessel wall. Variable resistances are modeled by combining a *tube law*, describing the mechanical response of the vessel wall to changes in the transmural pressure difference, and *Poiseuille law*, describing the fluid flow through the resistor. Following [21] and [29], we model deformable tubes as Starling resistors, which reflect the physiological high collapsibility of the venous segments when the transmural pressure becomes negative. For a generic resistance *R* we then write

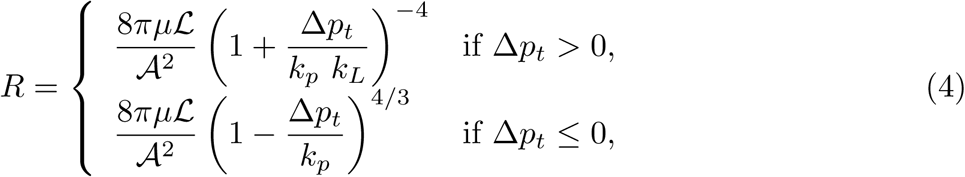

where,

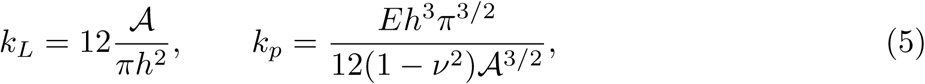

where *h, E* and *ν* are thickness, Young’s modulus and Poisson ratio of the vessel wall, respectively. The value of Δ*p*_*t*_ differs depending on the particular vascular segment under consideration, as specified in Table 2. In particular, in all circulation segments inside the eye (light blue shaded area in Fig 2) the pressure external to the vessel coincides with the IOP. On the other hand the translaminar segments of the central retinal artery and vein are subjected to an external radial compressive stress *P*_*LC*_ that originates within the lamina cribrosa as a consequence of the action of IOP, CSFp and scleral tension acting on it, as schematized in Fig 3 (left). In order to evaluate this compressive stress, we use the nonlinear elastic model proposed by some of the authors in [27], with the parameter choices reported in [21]. This three-dimensional numerical model is employed to compute *P*_*LC*_ for various values of IOP and CSFp forcing the lamina deformation. The results are reported with dots in Fig 3 (right) and have been fitted using a second order polynomial function, shown in the figure by the green surface. Such a surface is then employed to determine the external pressure acting on the translaminar vascular segments.

**Fig 3.**
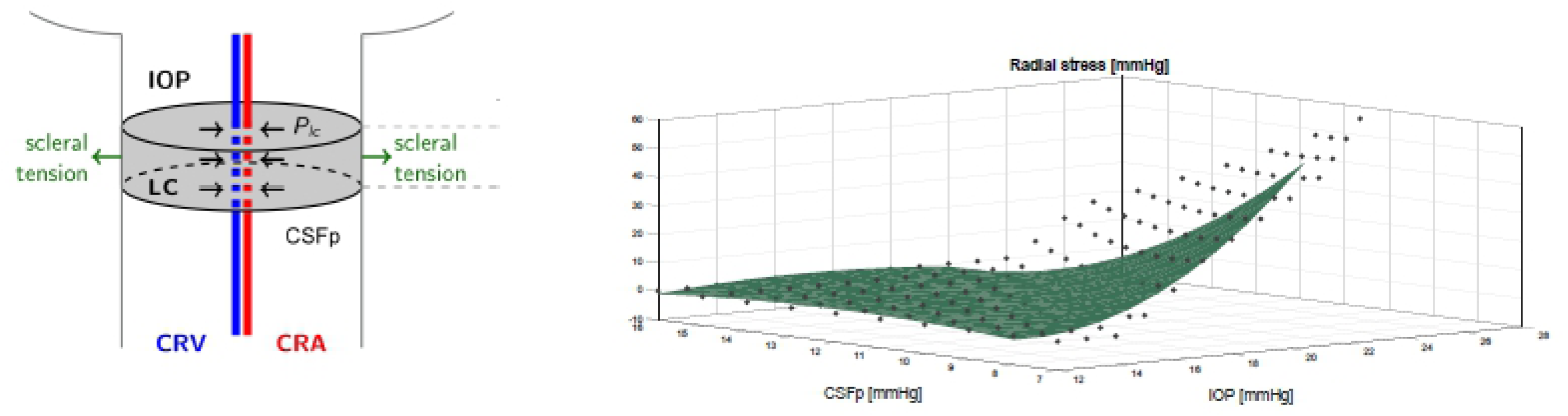
*Left:* Schematic representation of the radial compressive stress *P*_*LC*_ originating within the lamina cribrosa as a consequence of the action of IOP, CSFp and scleral tension acting on it. *P*_*LC*_ acts as external pressure on the translaminar segments of the central retinal artery and vein. *Right:* Magnitude of the radial compressive stress *P*_*LC*_ computed using the nonlinear elastic model described in [27] for various IOP and CSFp levels (black dots) and interpolated using using a second order polynomial function (green surface).

#### The aqueous humor circuit

The aqueous circuit used in this model was proposed by [22] and its electrical analogue is depicted in cyan in Fig 2. The value of IOP results from the balance between aqueous humor production and drainage.

Aqueous humor is produced at the ciliary body by a combination of passive and active mechanisms. The passive mechanism consists of ultrafiltration of fluid across the vessel wall in the microcirculation and produces a flux *J*_*uf*_ that is driven by trans-membrane differences in hydrostatic and oncotic pressures, the oncotic flux being mediated by a protein reflection coefficient *σ*_*p*_, see Eq (1). The passive production mechanism is accounted for in the model through the filtration block (resistance and current generator in parallel) between the capillary node in the ciliary circulation (with pressure *P*_*cl,c*_) and the node in the aqueous circuit (with pressure IOP), resulting in the flux *J*_*uf*_.

The active mechanism is due to secretion of ions in the non-pigmented epithelium, which creates a difference in ionic concentration that draws fluid into the eye via osmosis [23]. Following [22], we assume that the active secretion contributes with a constant flow rate, which is modeled in the electrical analogue of Fig 2 by the generator *J*_*secr*_. Thus, *J*_*secr*_ is a model input and is listed in Table 3.

**Table 3.**
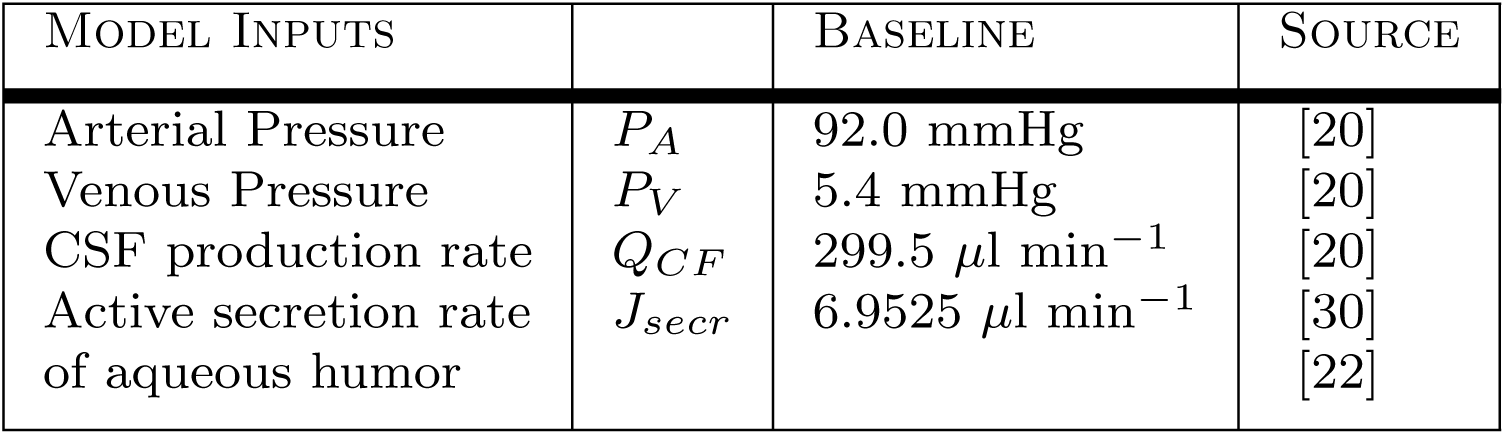
Summary of model inputs.

The drainage of aqueous humor from the eye is driven by mechanical pressure differences and occurs through two different pathways: the trabecular pathway (also known as conventional), resulting in the flow *J*_*tm*_ and the uveoscleral pathway (also known as unconventional), resulting in the flow *J*_*uv*_. Both are modeled as nonlinear resistors *R*_*tm*_ and *R*_*uv*_, as in [22].

Since the system is assumed to be at steady state, aqueous inflow and outflow must be equal to each other, leading to the balance equation *J*_*uf*_ + *J*_*secr*_ = *J*_*tm*_ + *J*_*uv*_. The full model of the eye results from coupling the circuits described above as shown in Fig 2. The coupling between the two circuits is truly two-ways: *(i)* the aqueous humor circuit influences the vascular circuit, since the IOP acts as an external pressure on the intraocular venules (see dashed cyan arrows in Fig 1), thereby influencing their resistances; *(ii)* the vascular circuit influences the aqueous humor circuit, since blood pressure in the ciliary body capillaries contributes to the production of aqueous humor.

### Coupling of the eye and brain models

The connections between the eyes and brain models are marked with special symbols in Fig 1. Red filled circles denote connections between arterial compartments; blue filled circles those between venous compartments. Finally, the cyan and grey arrows indicate the pressures (intraocular and extraocular, respectively) that act on both sides of the lamina cribrosa in the optic nerve head and on the vessels. More precisely:

- the node corresponding to the ophthalmic artery (OA) the eye model (whose pressure is denoted as *P*_*Eye,in*_ in Fig 1) is connected to the intracranial arteries in the brain model (whose pressure is denoted as *P*_*I*_) through an effective vascular resistance (*R*_*I,OA*_);
- the node corresponding to the cavernous sinus in the brain (whose pressure is denoted as *P*_*Eye,out*_) and that corresponding to the episcleral veins in the eye (whose pressure is denoted as *P*_*evp*_) are directly connected to the venous sinus in the brain model (whose pressure is denoted as *P*_*S*_);
- the lamina is acted upon by the IOP from the ocular side (cyan arrow) and by the CSFp in the subarachnoid space in the optic nerve, here assumed to be equal to the extraventricular CSFp (grey arrow).

Thank to this coupling, this work presents the first model capable of simulating in a self-consistent manner the fluid-dynamic eye-brain-body connections as the result of *(i)* the pressure drop between the central arteries (*P*_*A*_) and the central veins (*P*_*V*_); *(ii)* the active secretion of aqueous humor in the eye (*J*_*secr*_); and *(iii)* the production of CSF in the brain (*Q*_*CF*_), as summarized in Table 3.

### Simulation of microgravity conditions

In space, the human body experiences a series of changes in external conditions with respect to earth. The most obvious one is the lack of gravity that has an influence on the redistribution of fluids within the body. In particular, a significant fluid shift has been observed from the lower to the upper body and head. This is known to produce an increase in intracranial pressure and also a variation in the blood colloid osmotic pressure in the brain [5, 31]. No data exist on the response of a healthy blood-brain barrier to microgravity conditions. However, [20] suggested that such a barrier could be weakened by gravity unloading and showed, with the lumped parameter model of the brain that we also have adopted, that this is a relevant ingredient to reproduce the observed increase in ICP.

In the present paper, microgravity conditions are simulated following the same approach as in [20], namely: *(i)* we impose zero hydrostatic pressure distribution; *(ii)* we set the central venous pressure to zero; *(iii)* we decrease blood colloid osmotic pressure as a consequence of fluid shift towards the head; *(iv)* we weaken the blood-brain barrier by modifying the Starling-Landis equation coefficients (see Eq (1)). In particular, we increase the filtration coefficient *K*_*CB*_ between capillaries in the brain and the brain tissue and correspondingly decrease the reflection coefficient *σ*_*CB*_ as suggested by [20].

We also consider long head-down tilt (LHDT) conditions that are the most common earth clinical model used to mimic the effects of microgravity on body fluids shift. In particular, HDT is a ground-based experimental procedure, which simulates the effects of microgravity on the cardiovascular system: participants lay head down on a surface inclined by an angle *θ* = 6 degrees, with respect to a horizontal plane, and experience a shift of cephalic fluid similar to that occurring in microgravity. The position change affects the hydrostatic pressure distribution in the human body, which is equal to *ρgx* sin *θ*, with *x* being an axis running along the body and centered in the right atrium, *ρ* is fluid density and *g* is the gravitational acceleration. This means that the pressure variation between the central and intracranial compartments in the model can be written as *G* sin *θ*, with *G* = *ρgH*, and *H* is the distance between the right atrium and the base of the brain. We use our model to simulate LHDT, by changing both the gravitational direction and the blood oncotic pressure in the brain.

Summarizing, we simulate: *(i)* LHDT; *(ii)* microgravity environment with an intact blood/brain barrier (M0); and *(iii)* microgravity environment with a weakened blood/brain barrier (M1, M2), imposing the condition reported in Table 4.

**Table 4.** Conditions for simulated microgravity environment, as in [20].

|  | SIMULATION CONDITIONS |  |
| --- | --- | --- |
| LHDT | $P_{A,LHDT} = P_A + G \sin \theta$<br>$K_{CB,LHDT} = 0.0665 \text{ ml mmHg}^{-1} \text{ min}^{-1}$ | $P_{V,LHDT} = P_V + G \sin \theta$<br>$\sigma_{CB,LHDT} = 1$ |
| M0 | $P_{A,M0} = P_A$<br>$K_{CB,M0} = 0.0665 \text{ ml mmHg}^{-1} \text{ min}^{-1}$ | $P_{V,M0} = 0$<br>$\sigma_{CB,M0} = 1$ |
| M1 | $P_{A,M1} = P_A$<br>$K_{CB,M1} = 0.1164 \text{ ml mmHg}^{-1} \text{ min}^{-1}$ | $P_{V,M1} = 0$<br>$\sigma_{CB,M1} = 0.5714$ |
| M2 | $P_{A,M2} = P_A$<br>$K_{CB,M2} = 0.1330 \text{ ml mmHg}^{-1} \text{ min}^{-1}$ | $P_{V,M2} = 0$<br>$\sigma_{CB,M2} = 0.5$ |

In all four cases, model simulations are performed for decreasing values of the blood oncotic pressure, corresponding to different degrees of fluid shift towards the upper part of the body. Specifically, denoting by *π*_*c*_ the blood oncotic pressure, the oncotic pressure differences involved in the ultrafiltration of aqueous humor in the eye and CSF in the brain can be written as Δ*π*_*p*_ = *π*_*c*_ *− π*_*ah*_ and Δ*π*_*CB*_ = *π*_*c*_ *− π*_*B*_, respectively. Since *π*_*ah*_ *≈* 0 mmHg [23, 30], a reasonable baseline value for *π*_*c*_ is 25 mmHg. Simulations are performed for *π*_*c*_ ∈ (18.5, 25) mmHg. Decreases of 3.3 mmHg and 6.3 mmHg have been associated with LHDT and microgravity conditions, respectively, leading to *π*_*c*_ values of mmHg and 18.7 mmHg, respectively [20], which are indeed included in the simulated interval.

### Solution strategy

We implemented the circuit shown in Fig 1 in OpenModelica [32], which is a free, open-source modeling and simulation environment. In particular, we used the analogue electric components environment, which allows the user to draw a circuit and specify analytical expressions for nonlinear components. The circuit is automatically solved by the software. The graphic-oriented design of the circuit allows the user to easily modify the network and to include a large number of components.

### Model Calibration

The main goal of the model calibration is to select the values for all model parameters in such a way that, at baseline, the model predictions are in agreement with values reported in the literature. Since the choice of model parameters is one of the most delicate and important steps in devising a mathematical model, we report here the main rationale behind our choices. Whenever possible we adopted, the same parameter values as those reported in the literature, as summarized in Table 5. Unfortunately, though, some parameters pertaining to blood flow resistances were not readily available in the literature and needed to be calibrated starting from reference values of flow rates and pressures. In the following, we denote reference values using overlined symbols.

**Table 5.** Summary of model parameters.

| MODEL PARAMETERS |  |  |
| --- | --- | --- |
| **BRAIN** |  |  |
| *Cerebral Blood Flow* |  |  |
| $R_I$ | $33.5 \cdot 10^{-3} \text{ mmHg min ml}^{-1}$ | [20] |
| $R_C$ | $12.7 \cdot 10^{-3} \text{ mmHg min ml}^{-1}$ | [20] |
| $R_S$ | $12.7 \cdot 10^{-3} \text{ mmHg min ml}^{-1}$ | [20] |
| *Cerebrospinal and Interstitial Fluid Flow* |  |  |
| $K_{CB}$ | $0.0665 \text{ ml mmHg}^{-1} \text{ min}^{-1}$ | [20] |
| $\sigma_{CB}$ | 1 | [20] |
| $\Delta\pi_{CB}$ | 21 mmHg | [20] |
| $R_{BT}$ | $1.55 \text{ mmHg min ml}^{-1}$ | [20] |
| $R_{FT}$ | $0.6678 \text{ mmHg min ml}^{-1}$ | [20] |
| $R_{FB}$ | $0.9023 \text{ mmHg min ml}^{-1}$ | [20] |
| $R_{TS}$ | $9.90 \text{ mmHg min ml}^{-1}$ | [20] |
| **EYE** |  |  |
| *Blood Flow* |  |  |
| $E_v$ | 495 mmHg | [21] |
| $h_v$ | $7.73 \cdot 10^{-4} \text{ cm}$ | this work |
| $\mu_v$ | $2.12 \cdot 10^{-5} \text{ mmHg} \cdot \text{s}$ | this work |
| $\nu_v$ | 0.49 | [21] |
| $\mathcal{L}_v$ | 0.80 cm | this work |
| $R_{cl,in}$ | $16.70 \text{ mmHg min ml}^{-1}$ | this work |
| $R_{cl,a1} = R_{cl,a2}$ | $105.26 \text{ mmHg min ml}^{-1}$ | this work |
| $R_{cl,c1} = R_{cl,c2}$ | $33.83 \text{ mmHg min ml}^{-1}$ | this work |
| $\mathcal{A}_{cl,v}$ | $2.61 \cdot 10^{-4} \text{ cm}^2$ | this work |
| $R_{cl,out}$ | $23.90 \text{ mmHg min ml}^{-1}$ | this work |
| $R_{cra,in}$ | $376.76 \text{ mmHg min ml}^{-1}$ | this work |
| $\mathcal{L}_{cra,1} = \mathcal{L}_{cra,2}$ | 0.44 cm | [21] |
| $\mathcal{L}_{cra,3}$ | 0.02 cm | [21] |
| $\mathcal{L}_{cra,4}$ | 0.1 cm | [21] |
| $\mathcal{D}_{cra}$ | $175 \cdot 10^{-4} \text{ cm}$ | [21] |
| $E_{cra}$ | 2250 mmHg | [21] |
| $h_{cra}$ | $4 \cdot 10^{-3} \text{ cm}$ | [21] |
| $\mu_{cra}$ | $2.25 \cdot 10^{-5} \text{ mmHg} \cdot \text{s}$ | [21] |
| $\nu_{cra}$ | 0.49 | [21] |
| $R_{r,a1} = R_{r,a2}$ | $101 \text{ mmHg min ml}^{-1}$ | this work |
| $R_{r,c1} = R_{r,c2}$ | $94.33 \text{ mmHg min ml}^{-1}$ | this work |
| $\mathcal{A}_{r,v}$ | $1.55 \cdot 10^{-4} \text{ cm}^2$ | this work |
| $\mathcal{L}_{crv,1}$ | 0.1 cm | [21] |
| $\mathcal{L}_{crv,2}$ | 0.02 cm | [21] |
| $\mathcal{L}_{crv,3} = \mathcal{L}_{crv,4}$ | 0.44 cm | [21] |
| $\mathcal{D}_{crv}$ | $238 \cdot 10^{-4} \text{ cm}$ | [21] |
| $E_{crv}$ | 4500 mmHg | [21] |
| $h_{crv}$ | $10.7 \cdot 10^{-4} \text{ cm}$ | [21] |
| $\mu_{crv}$ | $2.43 \cdot 10^{-5} \text{ mmHg} \cdot \text{s}$ | [21] |
| $\nu_{crv}$ | 0.49 | [21] |
| $R_{crv,out}$ | $247.25 \text{ mmHg min ml}^{-1}$ | this work |
| $R_{ch,in}$ | $6.04 \text{ mmHg min ml}^{-1}$ | this work |
| $R_{ch,a1} = R_{ch,a2}$ | $38.12 \text{ mmHg min ml}^{-1}$ | this work |
| $R_{ch,c1} = R_{ch,c2}$ | $12.25 \text{ mmHg min ml}^{-1}$ | this work |
| $\mathcal{A}_{ch,v}$ | $5.06 \cdot 10^{-4} \text{ cm}^2$ | this work |
| $R_{ch,out}$ | $8.66 \text{ mmHg min ml}^{-1}$ | this work |
| *Aqueous Humour* |  |  |
| $L_{in}$ | $0.3 \mu\text{l min}^{-1} \text{ mmHg}^{-1}$ | [30] |
| $\sigma_p$ | 1 | [30] |
| $\Delta\pi_p$ | 25 mmHg | [30] |
| $k_1$ | $0.4 \mu\text{l min}^{-1}$ | [23] |
| $k_2$ | 5 mmHg | [23] |
| $R_0$ | $2.2 \text{ mmHg min } \mu\text{l}^{-1}$ | [38] |
| $\kappa$ | $0.012 \text{ mmHg}^{-1}$ | [38] |
| **BODY-BRAIN-EYE COUPLING** |  |  |
| $R_A$ | $101.5 \text{ mmHg min ml}^{-1}$ | [20] |
| $R_V$ | $407.99 \text{ mmHg min ml}^{-1}$ | [20] |
| $R_{TV}$ | $52.38 \text{ mmHg min ml}^{-1}$ | [20] |
| $R_{I,OA}$ | $18.27 \text{ mmHg min ml}^{-1}$ | this work |

#### Reference values for blood flow rates in the eye

A reference value of 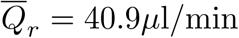 for retinal blood flow can be found in [21]. Similarly, a reference value of 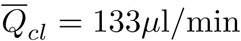 for the blood flow in the ciliary body can be found in [33]. The choice of a reference value for the choroidal blood flow is less obvious. Here we choose a value of 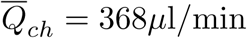, which is within the same range as the findings reported in [34] obtained using Magnetic Resonance Imaging, while yielding a percent ratio between retinal and choroidal blood flow of 10% to 90% as indicated in [11].

#### Reference values for blood pressures in the eye

A reference value of 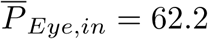 mmHg for blood pressure at the level of the ophthalmic artery can be found in [21]. This value follows from assuming that 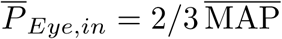, where 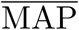 is the reference value for the mean arterial pressure defined as 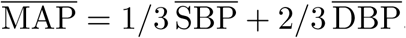, with 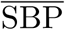 and 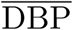 denoting the reference values for systolic and diastolic blood pressures, here set at 120 mmHg and 80 mmHg, respectively. A reference value of 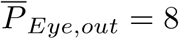 mmHg for blood pressure at the venous output of the eye can be found in [23]. References values for the pressure distribution inside the circuit for the retinal circulation can be found in Table 3 of [21]. Reference values of 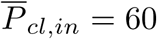 mmHg and 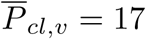 mmHg can be found in [23], whereas a reference value of 32 mmHg for the pressure between arterioles and capillaries can be found in [35] and [36]. Reference values for the pressures within the choroidal circulation have been set as to mirror those in the ciliary circulation, as in [23].

#### Reference values for vascular resistances in the eye

Given the reference values of flow rates and pressures, the values for vascular resistances can be found by means of Ohm’s law. The parameters involved in the calculation of the vascular resistances pertaining to CRA and CRV follow directly from the work by [21]. For what concerns the parameters involved in the resistances for retinal venules, we follow the same approach as in [21], where a dichotomous network model for the retinal circulation proposed in [37] was used to define the hierarchical architecture of the venous retinal segment, leading to the parameter values reported in Table 6. Owing to lack of data concerning ciliary and choroidal circulations, we assumed that all the elastic parameters are the same as those for the retina. For what concerns characteristic areas and lengths, we reasoned as follows. Let the blood volumes in the venous segments of retinal ciliary body and choroidal circulations be defined as 𝒱_*i,v*_ = *𝒜*_*i,v*_ *ℒ*_*i,v*_, for *i* = *r, cl, ch*. Let us assume that the ratio between blood volumes in the retinal and ciliary body (res. choroidal) venous segment is that same as the ratio between the corresponding flow rates, namely

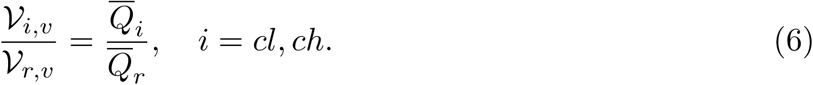

**Table 6.** Baseline values for pressures and flow rates under physiological condition.

| BASELINE VALUES |  |  |  |  |
| --- | --- | --- | --- | --- |
|  | PRESSURES | (mmHg) | FLOW RATES | (ml min<sup>-1</sup>) |
| BRAIN | $P_I$ | 81.9956 | $Q_{IC}$ | 1035 |
| | $P_C$ | 34.1543 | $Q_{CB}$ | 0.1291 |
| | $P_S$ | 7.8210 | $Q_{FT}$ | 0.3004 |
| | $P_B$ | 11.2125 | $Q_{FB}$ | $42.42 \cdot 10^{-3}$ |
| | $P_F$ | 11.2125 | $Q_{BT}$ | 0.1287 |
| | | | $Q_{TS}$ | 0.3220 |
| | | | $Q_{CS}$ | 1034.5 |
| EYE | $P_{cra,in}$ | 46.833 | $Q_r$ | $40.9 \cdot 10^{-3}$ |
| | $P_{cra,1}$ | 43.9041 | $Q_{ch}$ | $367.9 \cdot 10^{-3}$ |
| | $P_{cra,2}$ | 40.9753 | $Q_{cl}$ | $135.7 \cdot 10^{-3}$ |
| | $P_{cra,3}$ | 40.8465 | $J_{uf}$ | $-3.7 \cdot 10^{-3}$ |
| | $P_{cra,4}$ | 40.2141 | $J_{ah}$ | $3.3 \cdot 10^{-3}$ |
| | $P_{r,a}$ | 36.0875 | $J_{uv}$ | $2.9983 \cdot 10^{-4}$ |
| | $P_{r,c}$ | 28.1062 | $J_{tm}$ | 0.003 |
| | $P_{r,v}$ | 22.1371 | | |
| | $P_{crv,1}$ | 20.0226 | | |
| | $P_{crv,2}$ | 19.8107 | | |
| | $P_{crv,3}$ | 19.7729 | | |
| | $P_{crv,4}$ | 18.8452 | | |
| | $P_{crv,out}$ | 17.9235 | | |
| | $P_{ch,in}$ | 60.003 | | |
| | $P_{ch,a}$ | 45.9777 | | |
| | $P_{ch,c}$ | 27.4443 | | |
| | $P_{ch,v}$ | 16.9714 | | |
| | $P_{ch,out}$ | 11.0069 | | |
| | $P_{cl,in}$ | 60.0213 | | |
| | $P_{cl,a}$ | 46.1248 | | |
| | $P_{cl,c}$ | 27.7616 | | |
| | $P_{cl,v}$ | 17.118 | | |
| | $P_{cl,out}$ | 11.0651 | | |
| | $P_{LC}$ | 0 | | |
| | $IOP$ | 14.9662 | | |
| | $P_{evp}$ | 7.82104 | | |
| BODY-BRAIN-EYE<br>COUPLING | $P_{Eye,in}$ | 62.2270 | $Q_A$ | 1035 |
| | $P_T$ | 11.0119 | $Q_{TV}$ | 0.1071 |
| | $P_{Eye,out}$ | 7.8210 | $Q_{CVS} = Q_{Eye,out}$ | 0.5475 |
| | $RLTp$ | 11.0119 | $Q_{SV}$ | 1035 |
| | | | $Q_{OA} = Q_{Eye,in}$ | 0.5407 |
| | | | $Q_V$ | 1035 |

Then it follows that

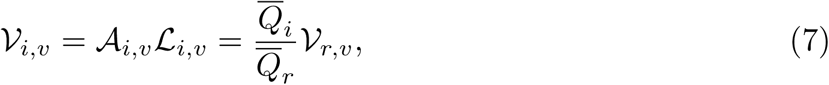

which implies that

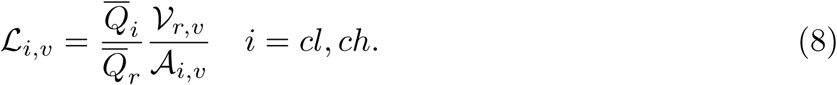

The value of *𝒜*_*i*_, *i* = *cl, ch*, is computed imposing that, when 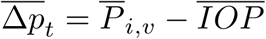, with 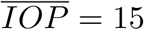 mmHg, the value of the resistance in the Starling model equals the value of 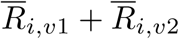 obtained using Ohm’s law.

The solution of the system obtained for the model parameters reported in Table 5 leads to the baseline distribution of pressures and flow rates reported in Table 6. A thorough validation of the model is the object of the next section.

### Model validation

The mathematical model presented here connects numerous blocks that have been developed independently by different authors. Thus, the validation of each block considered individually has been addressed in different articles. Specifically, for the circulation in the retina and in the central retinal vessels we refer to [21], for the circulation in the ciliary body and in the choroid we refer to [39] and [23], for the aqueous humor circulation we refer to [22] and for the cerebral circulation of blood, interstitial fluid and CSF we refer to [20]. The validation of the modeling assumptions concerning the connections between the various components has been presented at the 2018 Annual Meeting of the Association for Research in Vision and Ophthalmology [10]. The model proved capable of capturing a slight increase in IOP following an increase in blood pressure, consistently with the findings of the Rotterdam study [40], the Blue Mountains Eye population-based study [41] and the Beijing eye study [42]. The model also predicted a decrease in CSFp with blood pressure, consistently with the findings of Ren and Wang reported in [43]. In addition, the model confirmed that the pressure in the vortex veins before they exit the sclera is approximately equal to the IOP over a wide range of values, as reported by Bill in [44]. These validation results are quite impressive, considering that these studies were not used for model calibration. The successful validation of the model blocks considered separately [20–23, 39] and jointly [10] supports the utilization of the model to study the physiological connections between ocular and cerebral fluid flows.

## Results and Discussion

We now turn to the discussions of the results obtained by simulating microgravity conditions using the mathematical model presented in the previous sections. We consider the four different cases summarized in Table 4, namely LHDT, M0, M1 and M2.

Changes in the blood oncotic pressure modify both Δ*π*_*CB*_ and Δ*π*_*p*_, leading to changes in the filtration across the blood-brain barrier and the ciliary body, respectively, and, consequently, to changes in ICP, CSFp and IOP. This is a crucial ingredient to interpret the results of the model. We also note that the model is highly nonlinear and, in particular, the Starling resistors in the venous segments are likely to play an important role. Specifically, when the transmural pressure becomes negative the venous compartments collapse, giving rise to a significant increase in resistance, according to Eq (4).

Fig 4 shows IOP (cyan), ICP (grey) and the radial stress acting on *cra* and *crv* (green) as a function of the blood oncotic pressure *π*_*c*_ for the four cases of interest (LHDT, M0, M1 and M2). We recall that a reasonable baseline value for *π*_*c*_ on earth is *≈*25 mmHg, which may decrease to 21.7 mmHg in LHDT and to 18.7 mmHg in microgravity conditions. These reference values are indicated with dashed lines in the corresponding figures.

**Fig 4.**
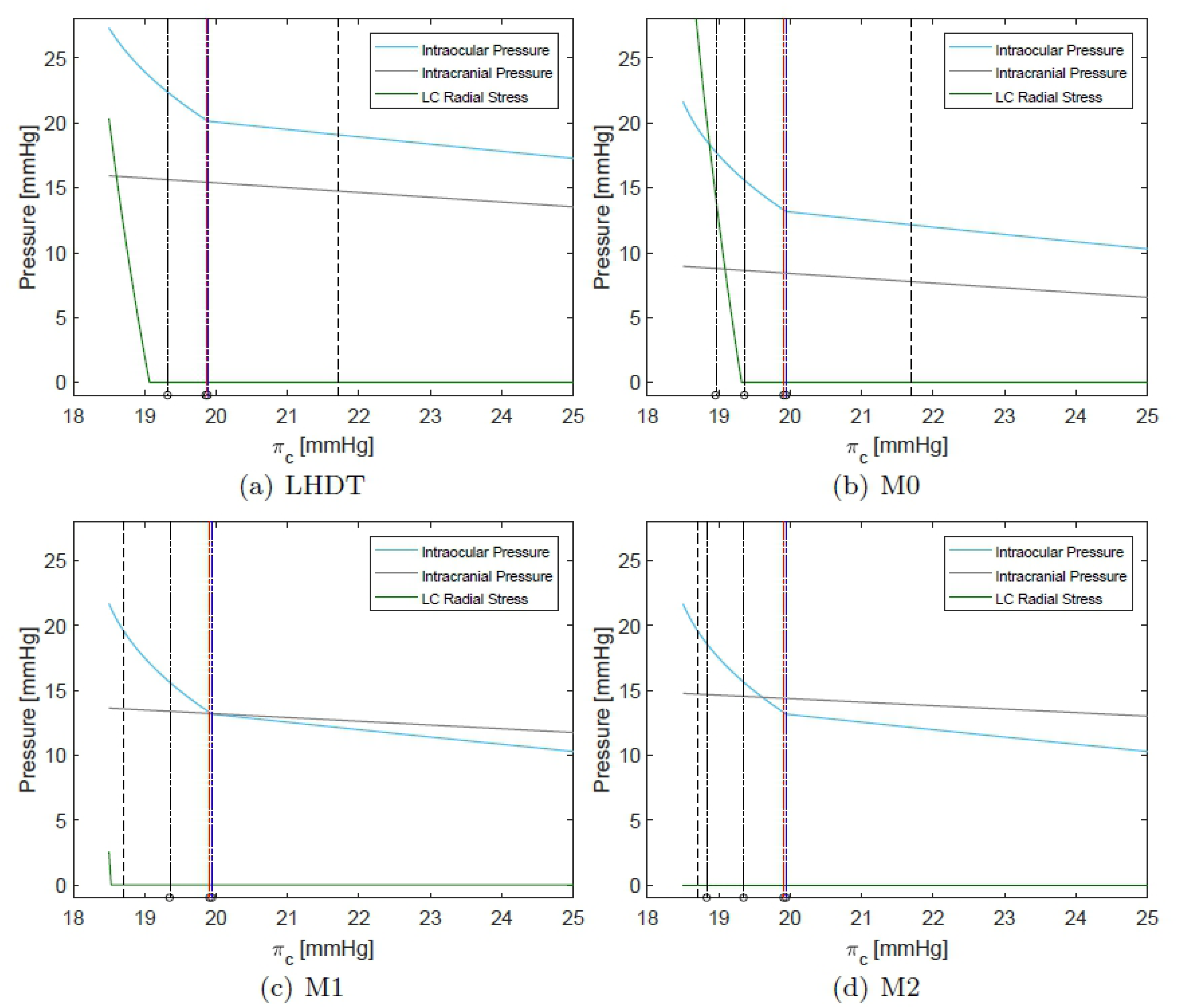
Intraocular, intracranial and compressive radial stress within the lamina cribrosa (LC) as a function of the blood oncotic pressure in capillaries *π*_*c*_. The physiological value of *π*_*c*_ on earth is *≈* 25 mmHg. The vertical dotted line indicates the value of *π*_*c*_ representative of each case. Dashed coloured vertical lines indicate the values of *π*_*c*_ at which each of compartment collapses (black for central retinal vein, red for choroidal venules and blue for ciliary venules).

In all cases reported in Fig 4, and in agreement with the results of [20], the intracranial pressure grows as *π*_*c*_ decreases and the increase is more marked when the weakening of the blood/brain barrier is accounted for (M1 and M2). The ICP variation is approximately linear with *π*_*c*_.

IOP also grows as the oncotic pressure difference decreases. In the Starling resistors, collapse of the flexible pipes depends on the transmural pressure that, in all cases but the prelaminar *crv*, is given by the difference between the intraluminal pressure and the IOP. Therefore, a growth in IOP can lead to vessel collapse. Since this is an important ingredient in the model, in Fig 4 we report with coloured dotted lines the values of *π*_*c*_ at which each of the compartments collapses. Throughout this section, we use consistently the following *color code* to identify different ocular blood compartments:

- black for retinal circulation;
- red for choroidal circulation; and
- blue for ciliary body circulation.

Thus, for example, black vertical lines in Fig 4 indicate the maximum value of *π*_*c*_ for which vessel collapse in the retinal circulation occurs. In all considered cases, the first compartment to collapse (for the largest values of *π*_*c*_) is that corresponding to the ciliary venules. Soon afterwards, and further decreasing *π*_*c*_, the choroidal venules and the postlaminar *crv* also collapse. Finally, the prelaminar *crv* collapses at a slightly smaller value of *π*_*c*_. This last event is triggered by a compressive radial stress acting on the vessel, produced by the lamina deformation. We note that all these events occur in a relatively narrow range of values of the blood oncotic pressure *π*_*c*_ and, approximately, at the same values of *π*_*c*_ in all considered cases. In the case of LHDT the collapse is slightly delayed.

We now analyze in detail the variation of the IOP with *π*_*c*_ shown in Fig 4 and discuss its implications for blood fluxes in the various ocular compartments. For the sake of clarity we will consider first the case of LHDT and then compare it with the microgravity cases (M0, M1 and M2). In Fig 4(a) the IOP varies almost linearly with *π*_*c*_, in the range 20 mmHg ⪅ *π*_*c*_ *≤*25 mmHg and, for *π*_*c*_ ⪅ 20 mmHg, the dependency becomes markedly nonlinear. This change in behavior is related to the collapse of ciliary venules that occurs when the transmural pressure in the ciliary venules becomes negative. The collapse of the venules produces a pressure increase in the ciliary body capillaries and arterioles. This increases the aqueous production rate and, in turn, makes the IOP grow. Finally, we note that the radial stress in the lamina cribriosa remains almost constant over a wide range of values of *π*_*c*_ and then sharply increases.

The fluxes in the retina, choroid and ciliary body circulation systems, normalized with their respective baseline values, are reported in Fig 5. The figure shows that, as *π*_*c*_ decreases, the ciliary and choroidal fluxes decrease almost linearly until vessel collapse occurs, and then do so much more rapidly. Interestingly, a different behavior is predicted by the model for the retina. Over a wide range of values of *π*_*c*_ and before venules collapse happens, the flux in the retina remains almost constant. This suggests that the particular architecture of the retinal vasculature provides a sort of mechanical (i.e. purely passive) blood flow regulation in response to IOP changes. This behavior was already observed in [21], who proposed a model of the retinal circulation alone. It is interesting to find that this feature is maintained by the full coupled, interconnected model. When venules collapse in the ciliary body and choroidal circulation, the flux in the retina initially grows. This is due to the fact that the retinal circuit is arranged in parallel to the other vascular beds and a decrease in flux in the ciliary body and choroid is partially compensated by an increase in the retinal blood flow. Once the *crv* also collapses, blood flow in the retina starts to drop significantly.

**Fig 5.**
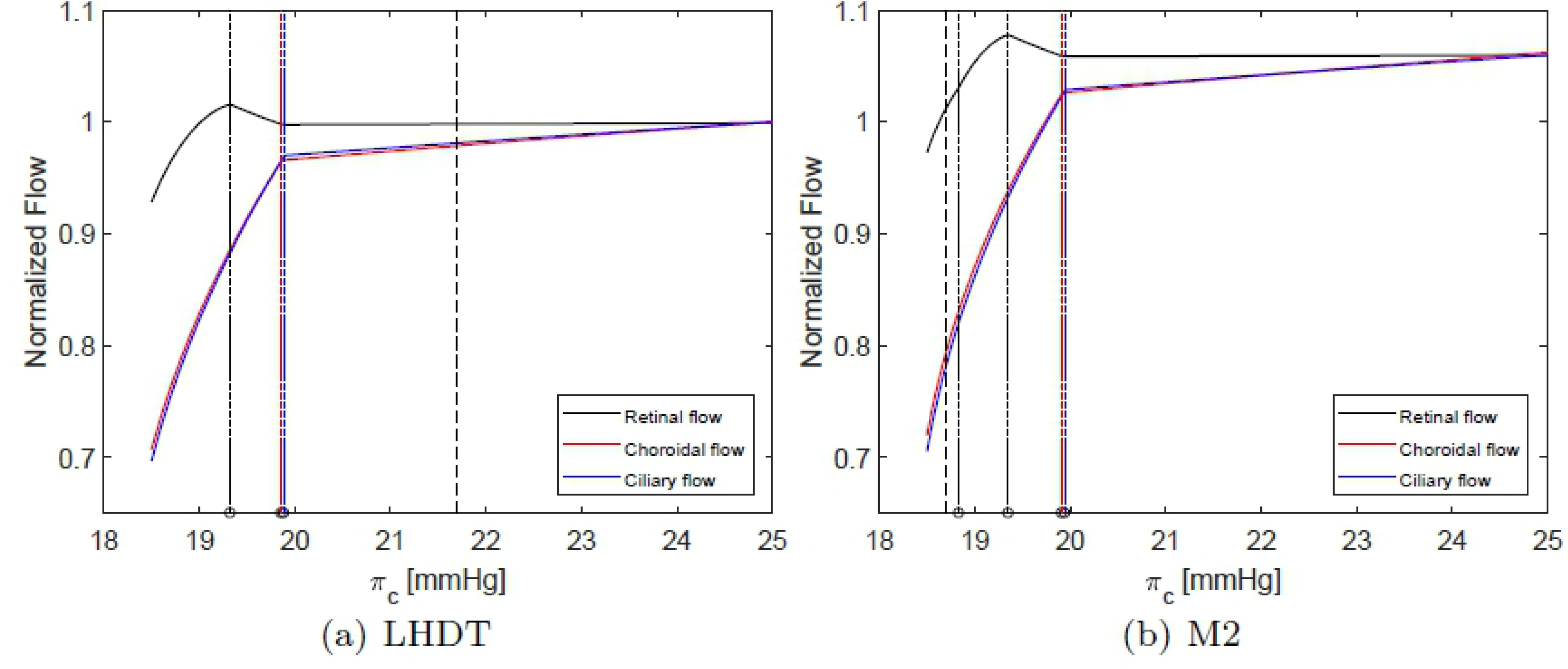
Normalized fluxes in the retina, choroid and ciliary body as a function of the blood oncotic pressure in capillaries *π*_*c*_. All fluxes are normalized with their respective baseline value. The vertical dashed line indicates the value of *π*_*c*_ representative of each case. Dotted colored vertical lines indicate the values of *π*_*c*_ at which each of compartment collapses (black for central retinal vein, red for choroidal venules and blue for ciliary venules).

Figs 6 and 7 show the dependency of the transmural pressure in various compartments on the blood oncotic pressure *π*_*c*_. All the plots can be interpreted in the light of the comments above and provide a detailed picture of pressure changes in the ocular circulation as a response to the LHDT experiment.

**Fig 6.**
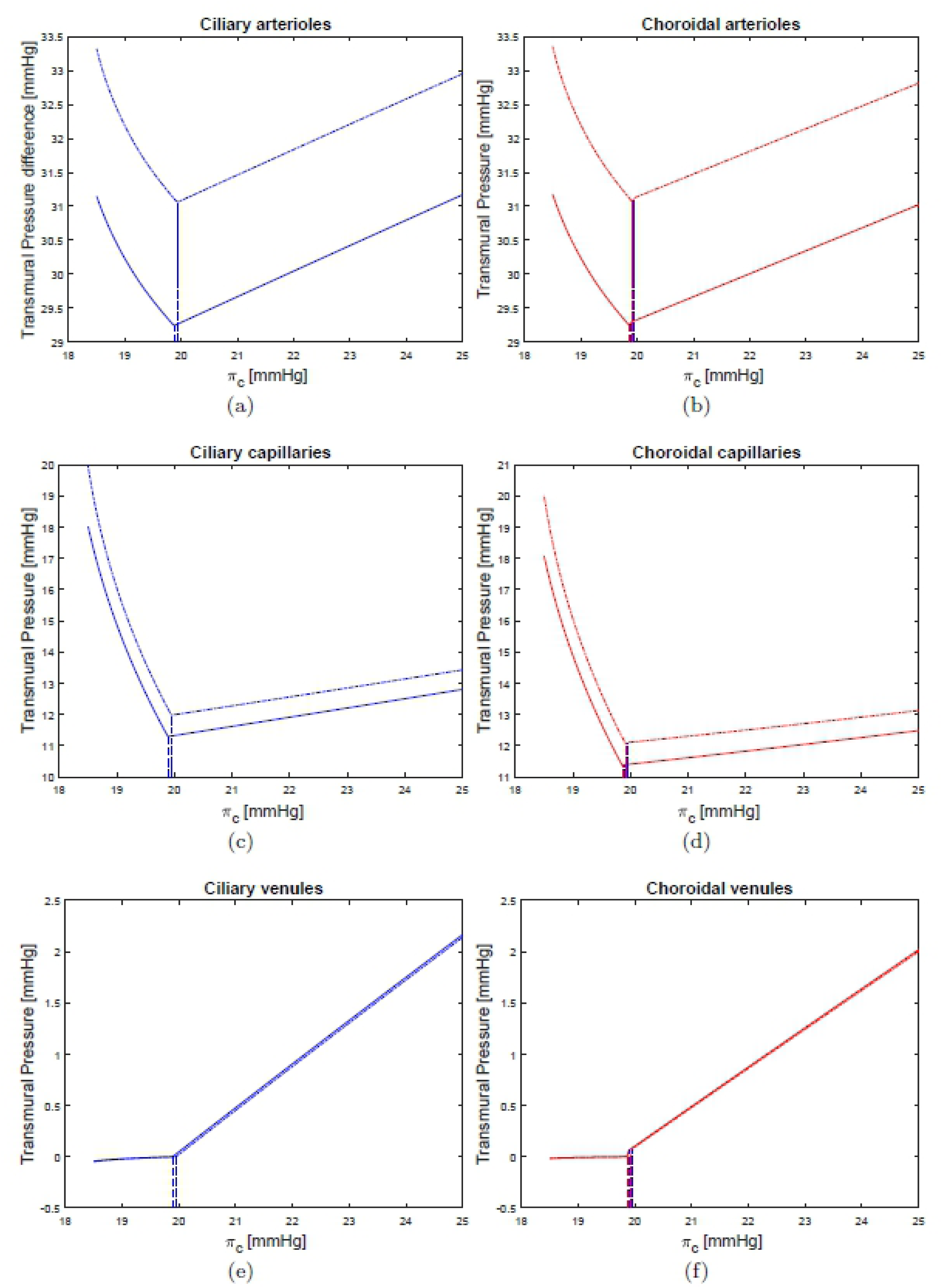
Transmural pressures in various compartments of the ciliary and choroidal circulations. LHDT solid line, M2 dash-dotted line.

**Fig 7.**
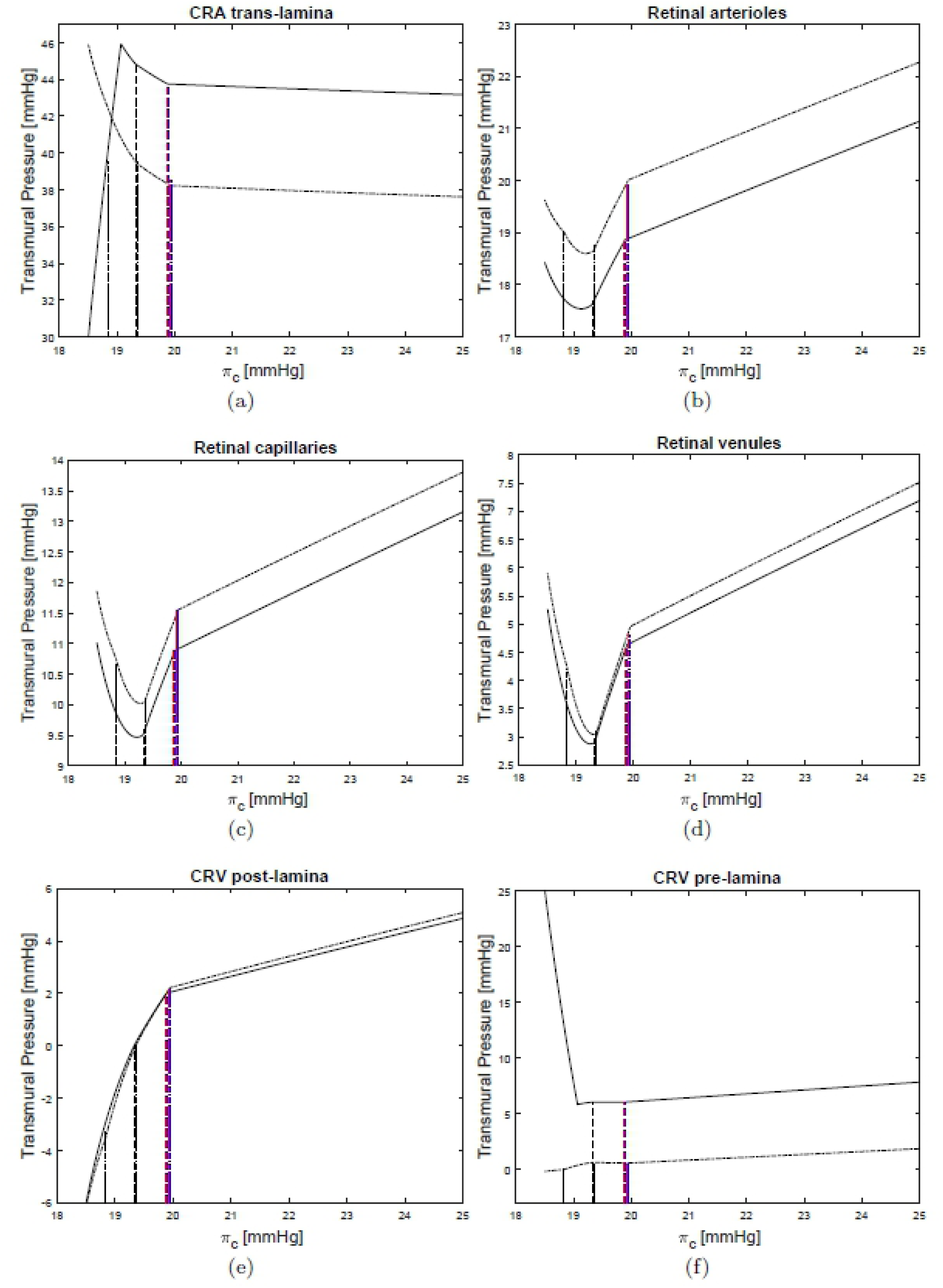
Transmural pressures in various compartments of the retinal circulation. LHDT solid line, M2 dash-dotted line.

Finally, the LHDT case is compared with the simulations of microgravity conditions. Fig 4(b-d) shows that in microgravity conditions the ICP is smaller than in the case of LHDT, when the effect of blood/brain barrier disruption is not accounted for. ICP increases as the filtration and reflection coefficients are changed. These results are in agreement with the findings of [20]. The IOP predicted by our model is invariably lower in the cases of microgravity than of LHDT, even if it is larger than in physiological earth conditions. The effect of the blood/brain barrier integrity is far less important for the IOP than it is for the intracranial pressure. It is noted that in Figs 6 and 7 we only report the curves corresponding to the case M2, since they are almost coincident with those relative to the cases M0 and M1. Concerning blood fluxes, the dependency on *π*_*c*_ is similar to the LHDT case. However, the flux can be higher than in baseline earth conditions, especially in the retina. This is essentially due to the lowering of the central venous pressure.

## Conclusion

In this paper, a lumped parameter model of fluid flow in the eyes and brain is proposed to study the flow of blood, interstitial fluid, CSF and aqueous humor. Despite its many limitations, including the fact that time-dependence and vascular regulation are not accounted for, the model proved capable of capturing, both qualitatively and quantitatively, the relationships between IOP, CSFp and blood pressure reported in major clinical and population-based studies. In addition, the model is able to reproduce the seminal result regarding choroidal venous pressure by [44], thereby suggesting that it may prove instrumental to study blood outflow through the vortex veins, which is thought to play a very important role in many pathologies but remains very difficult to measure.

The model has been utilized to investigate changes in flow and pressure distributions associated with long term exposure to microgravity conditions. In this respect our model is complementary to the one proposed in [6], which was focused on the short term effects induced by acute gravitational variations. Simulation results point at a purely mechanical feedback mechanism due to the collapsibility of the retinal venules exposed to IOP that would help maintain a relatively constant level of blood flow trough the retina despite microgravity-induced changes in pressure. Thus, a certain level of regulation is achieved in the retina even though active regulatory mechanisms are not included in the model. Interestingly, a similar mechanism is not present in the choroid and ciliary body due to their different vascular architectures. We hypothesize that this could be one of the reasons why SANS affects the choroid and the optic nerve more than the retina [1]. It is important to notice that various conditions during space flight are thought to alter vascular regulatory capabilities, particularly the increased amount of CO_2_ in the air. With this in mind, a purely passive regulatory mechanism, as that predicted by the model for the retinal circulation, might turn out to be of great importance to avoid retinal damages in space.

Thanks to its ability to self-consistently compute IOP and CSFp given a certain level of blood pressure, the model developed in this paper might also bear a great relevance for other pathologies beyond SANS, most importantly glaucoma. In addition, the ability to simulate the flow conditions in the venous segments of the eye is a major contribution in ophthalmology, since the veins are known to play a very important role in ocular physiology but are very difficult to measure in vivo.

We conclude by emphasizing that the model indeed includes a large number of parameters, which required careful calibration. Despite the very good results obtained on the model validation, it would be important to perform a sensitivity analysis on the paramater space to better assess the robustness of the model and direct its future extensions.

## Acknowledgments

This work has been partially supported by the award NSF DMS-1853222/1853303, the Chair Gutenberg funds of the Cercle Gutenberg (France) and the Labex IRMIA and the IdEx Unistra (University of Strasbourg, France).

